# Methodologic Issues in Doubly Labeled Water Measurements of Energy Expenditure During Very Low-Carbohydrate Diets

**DOI:** 10.1101/403931

**Authors:** Kevin D. Hall, Juen Guo, Kong Y. Chen, Rudolph L. Leibel, Marc L. Reitman, Michael Rosenbaum, Steven R. Smith, Eric Ravussin

## Abstract

**Background:** Very low-carbohydrate diets have been reported to substantially increase human energy expenditure as measured by doubly labeled water (DLW) but not by respiratory chambers. Do the DLW data reflect true physiological differences that are undetected by respiratory chambers? Alternatively, are the apparent DLW energy expenditure a consequence of failure to fully account for respiratory quotient (RQ) differences between diets?

**Objective:** To examine energy expenditure differences between diets varying drastically in carbohydrate and to quantitatively compare DLW data with respiratory chamber and body composition measurements within an energy balance framework.

**Design:** DLW measurements were obtained during the final two weeks of month-long baseline (BD; 50% carbohydrate, 35% fat, 15% protein) and isocaloric ketogenic diets (KD; 5% carbohydrate, 80% fat, 15% protein) in 17 men with BMI 25-35 kg/m^2^. Subjects resided 2d/week in respiratory chambers to measure energy expenditure (EE_chamber_). DLW expenditure was calculated using chamber-determined respiratory quotients (RQ) either unadjusted (EE_DLW_) or adjusted (EE_DLWΔRQ_) for net energy imbalance using diet-specific coefficients. Accelerometers measured physical activity. Body composition changes were measured by dual-energy X-ray absorptiometry which were combined with energy intake measurements to calculate energy expenditure by balance (EE_bal_).

**Results:** After transitioning from BD to KD, neither EE_chamber_ nor EE_bal_ were significantly changed (∆EE_chamber_=24±30 kcal/d; p=0.43 and ∆EE_bal_=-141±118 kcal/d; p=0.25). Similarly, physical activity (−5.1±4.8%; p=0.3) and exercise efficiency (−1.6±2.4%; p=0.52) were not significantly changed. However, EE_DLW_ was 209±83 kcal/d higher during the KD (p=0.023) but was not significantly increased when adjusted for energy balance (EE_DLWΔRQ_ =139±89 kcal/d; p=0.14). After removing 2 outliers whose EE_DLW_ were incompatible with other data, EE_DLW_ and EE_DLW∆RQ_ were marginally increased during the KD by 126±62 kcal/d (p=0.063) and 46±65 kcal/d (p=0.49), respectively.

**Conclusions:** DLW calculations failing to account for diet-specific energy imbalance effects on RQ erroneously suggest that very low carbohydrate diets substantially increase energy expenditure.

## Introduction

There is a great deal of interest in whether very low-carbohydrate diets offer a “metabolic advantage” for weight loss by increasing total energy expenditure (1, 2) – a phenomenon predicted by the carbohydrate-insulin model of obesity (3, 4). In support of this concept, a highly-cited controlled feeding study in humans using the doubly labeled water (DLW) method found a substantial increase in average daily energy expenditure during a very low-carbohydrate diet compared with an isocaloric high-carbohydrate diet (5). However, such results are inconsistent with several controlled feeding studies employing respiratory chambers that failed to detect important energy expenditure differences between isocaloric diets varying in carbohydrate content (6-8).

Our recent controlled feeding inpatient study was the first to compare daily energy expenditure between isocaloric diets widely varying in carbohydrate content using both respiratory chambers and DLW (9). Our primary result was a small but statistically significant increase daily energy expenditure as measured by respiratory chamber (EE_chamber_) of 57 ± 13 kcal/d (P = 0.0004) after 17 men (25<BMI<35 kg/m^2^) transitioned from a one-month inpatient run-in period consuming a moderate-carbohydrate baseline diet (BD) (50% carbohydrate, 35% fat, 15% protein) to a subsequent month-long inpatient period consuming an isocaloric ketogenic diet (KD) (5% carbohydrate, 80% fat, 15% protein). An exploratory aim of our study was to measure changes in average daily energy expenditure by the DLW method (EE_DLW_) and we reported a more substantial increase of 151 ± 63 kcal/d (P = 0.03 uncorrected for multiple comparisons) following the KD (9).

Some investigators who promote the carbohydrate-insulin model of obesity have discounted these EE_chamber_ results and have reinterpreted our exploratory EE_DLW_ data as underestimating the true physiological effect of very low carbohydrate diets which have been suggested to increase energy expenditure by 200-300 kcal/d (10, 11). However, there are methodological reasons why the energy expenditure calculated by the DLW method is impacted by dietary carbohydrate content that do not reflect true physiological differences. Specifically, the DLW method requires an estimate of the overall metabolic fuel utilization of the body as quantified by the average daily respiratory quotient (RQ). While it is widely known that RQ depends on the composition of the diet as well as the overall state of energy balance, it is less well appreciated that the energy imbalance adjustment of RQ also depends on dietary carbohydrate (12) and such adjustments are rarely employed in DLW calculations.

Here, we reanalyzed our DLW data taking account of the diet-specific energy imbalance adjustments of RQ used in the DLW method and highlight some key challenges to interpreting DLW data when comparing energy expenditure differences between diets varying in carbohydrate.

## Methods

Details of the study and its approval by the Institutional Review Boards were reported previously (9). Briefly, 17 men with BMI between 25-35 kg/m^2^ provided informed consent and were admitted as inpatients to metabolic wards where they consumed a standard baseline diet (BD) composed of 50% energy from carbohydrate, 35% fat, and 15% protein for 4 weeks immediately followed by 4 weeks of an isocaloric very low-carbohydrate, ketogenic diet (KD) composed of 5% carbohydrate, 80% fat, 15% protein. Body weight and height were measured to the nearest 0.1 kg and 0.1 cm, respectively, with subjects wearing a hospital gown and undergarments and following an overnight fast. Body fat was measured using dual-energy X-ray absorptiometry (DXA) scanners (Lunar iDXA, GE Healthcare, Madison, WI, USA).

Subjects spent two consecutive days each week residing in respiratory chambers to measure energy expenditure (EE_chamber_). As described previously (9), during the BD period, the daily energy expenditure was calculated as follows: 
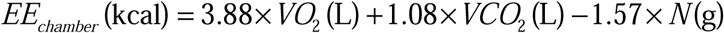
where *VO*_2_ and *VCO*_2_ were the volumes of oxygen consumed and carbon dioxide produced, respectively, and *N* was the 24hr urinary nitrogen excretion measured by chemiluminescence (Antek MultiTek Analyzer, PAC, Houston, TX). During the KD period, the equations were adjusted to account for 24-hour urinary ketone excretion, *K_excr_*: 
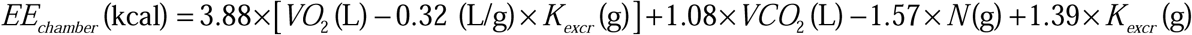

Energy efficiency of physical activity was measured in the respiratory chamber with subjects exercising at a constant, self-selected, level of moderate-intensity cycle ergometry.

Energy expenditure was calculated by energy balance (EE_bal_) using the daily metabolizable energy intake (EI) and the measured rates of change of the body energy storage pools determined from measurements of fat mass (FM) and fat-free mass (FFM) by DXA at the beginning and end of each two-week BD and KD period coincident with the DLW measurements as follows: 
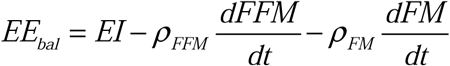
where ρ_FM_ = 9300 kcal/kg is the energy density of body fat mass, ρ_FFM_ = 1100 kcal/kg is the energy density of fat-free mass.

Energy expenditure was measured by DLW during the final two weeks of the BD and KD periods to allow sufficient time for fluid shifts as subjects adjusted to each diet. Subjects drank from a stock solution of ^2^H_2_O and H_2_^18^O water in which 1 g of ^2^H_2_O (99.99% enrichment) was mixed with 19 g of H_2_^18^O (10% enrichment). An aliquot of the stock solution was saved for dilution to be analyzed along with each set of urine samples. The water was weighed to the nearest 0.1 g into the dosing container. The prescribed dose was 1.0 g per kg body weight and the actual dose amounts were entered in the dose log. Spot urine samples were collected daily. Isotopic enrichments of urine samples were measured by isotope ratio mass spectrometry. The average CO_2_ production rate (rCO_2_) was corrected for previously administered isotope doses (13), can be estimated from the rate constants describing the exponential disappearances of the labeled ^18^O and deuterated water isotopes (*k_O_* and *k_D_*) in repeated spot urine samples collected over several days. We used the parameters of Racette et al. (14) with the weighted dilution space calculation, *R_dil_*, proposed by Speakman (15): 
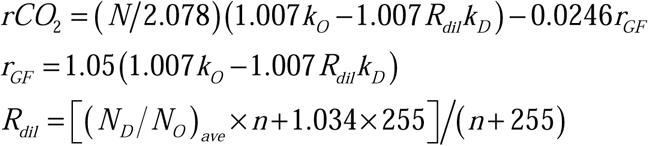
where *r_GF_* accounts for the fractionation of the isotopes and (*N_D_ / N_O_*)_*ave*_ is the mean of the *N_D_ / N_O_* values from the *n*=17 subjects.

In our previous report (9), we used the 24hr respiratory quotient, RQ, measured during the respiratory chamber stays to calculate energy expenditure (EE_DLW_) during the baseline period was calculated as: 
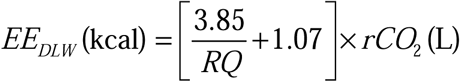

During the KD period, EE_DLW_ was calculated as: 
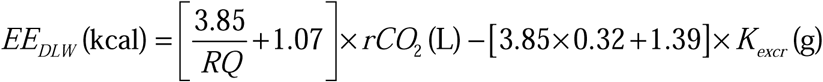

However, because the energy expended outside the chamber was significantly greater than inside the chamber (9), the chamber RQ values did not accurately represent the overall RQ values during the DLW period. In other words, the relative magnitude of energy imbalance (EB) during the DLW period was different than the energy imbalance (EI-EE_chamber_) measured during the chamber stays. Therefore, we used the overall energy imbalance during the DLW period, defined by the rate of change of body energy stores, to adjust the chamber RQ measurements by an amount ∆RQ as described in the Online Supplemental Materials: 
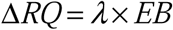
where λ = 5.28×10^-5^ d/kcal for the BD and λ = 9.17×10^-6^ d/kcal for the KD.

Importantly, the value of λ is affected by dietary carbohydrate content, and, consequently, the energy equivalence of CO_2_ produced in the DLW calculations differs between diets varying in carbohydrate for the same degree of energy imbalance.

Thus, the ΔRQ adjustment for each individual’s diet-specific state of energy imbalance results in the following calculation of DLW energy expenditure (EE_DLWΔRQ_) during the baseline period: 
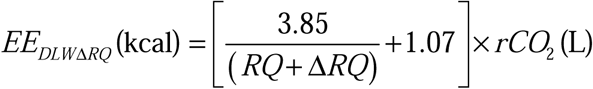

During the KD period, EE_DLWΔRQ_ was calculated as: 
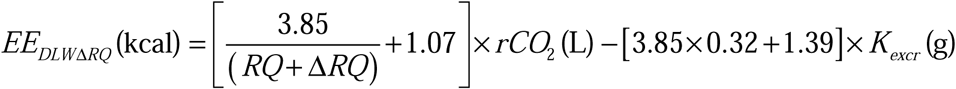

As opposed to our previous study (9) that reported results during the entire six-week period when EI was held constant (i.e. the last two weeks of the BD phase and the entirety of the KD phase), we now report results based upon data obtained only during the two-week DLW phase of both BD and KD periods. In addition to allowing direct comparison with the coincident DLW measures, focusing on the last two weeks of each diet allowed for two weeks to adapt to the BD from the usual diet as well as to the KD from BD.

Statistical analysis was performed using a paired, two-sided t-test with significance declared at the p<0.05 threshold. The data are reported as mean±SE.

## Results

During the final two weeks of the BD, EI was 2738±107 kcal/d which was significantly higher than EE_chamber_ = 2626±104 kcal/d (p<0.0001). EE_DLW_ was 2964±126 kcal/d and significantly higher than EI (p=0.011). After adjusting for the energy imbalance, EE_DLW∆RQ_ = 3045±135 kcal/d was significantly greater than EE_DLW_ (p=0.0003). Energy expenditure calculated using EI and the changes in body energy stores was EE_bal_ = 3136±171 kcal/d and was not significantly different from EE_DLW_ (p=0.23) or EE_DLW∆RQ_ (p=0.47). Compared to EE_chamber_, EE_DLW_ was 338±77 kcal/d higher (p=0.0005), EE_DLW∆RQ_ was 419±76 kcal/d higher (p<0.0001), and EEbal was 509±100 kcal/d higher (p<0.0001) indicating that subjects expended significantly more energy outside the chamber during the BD phase.

During the final two weeks of the KD phase, EI was 2730±110 kcal/d which, by design, was not significantly different from the BD phase (p=0.16). Whereas we previously reported a transient increase in EE_chamber_ during the first two weeks after introducing the KD (9), neither EE_chamber_ (2650±89 kcal/d; p=0.43) nor EE_bal_ (2995±160 kcal/d; p=0.25) were significantly different during the last two weeks of the KD compared to the last two weeks of the BD coincident with the DLW measurement periods (Figure 1A). Likewise, physical activity measured using an accelerometer mounted on the hip was not significantly different (KD relative to BD, −5.1±4.8%; p=0.3); and energy efficiency of physical activity measured in the respiratory chamber with subjects exercising at a constant level of moderate-intensity cycle ergometry was not significantly different (−1.6±2.4%; p=0.52) between the BD and KD phases.

**Figure 1.**
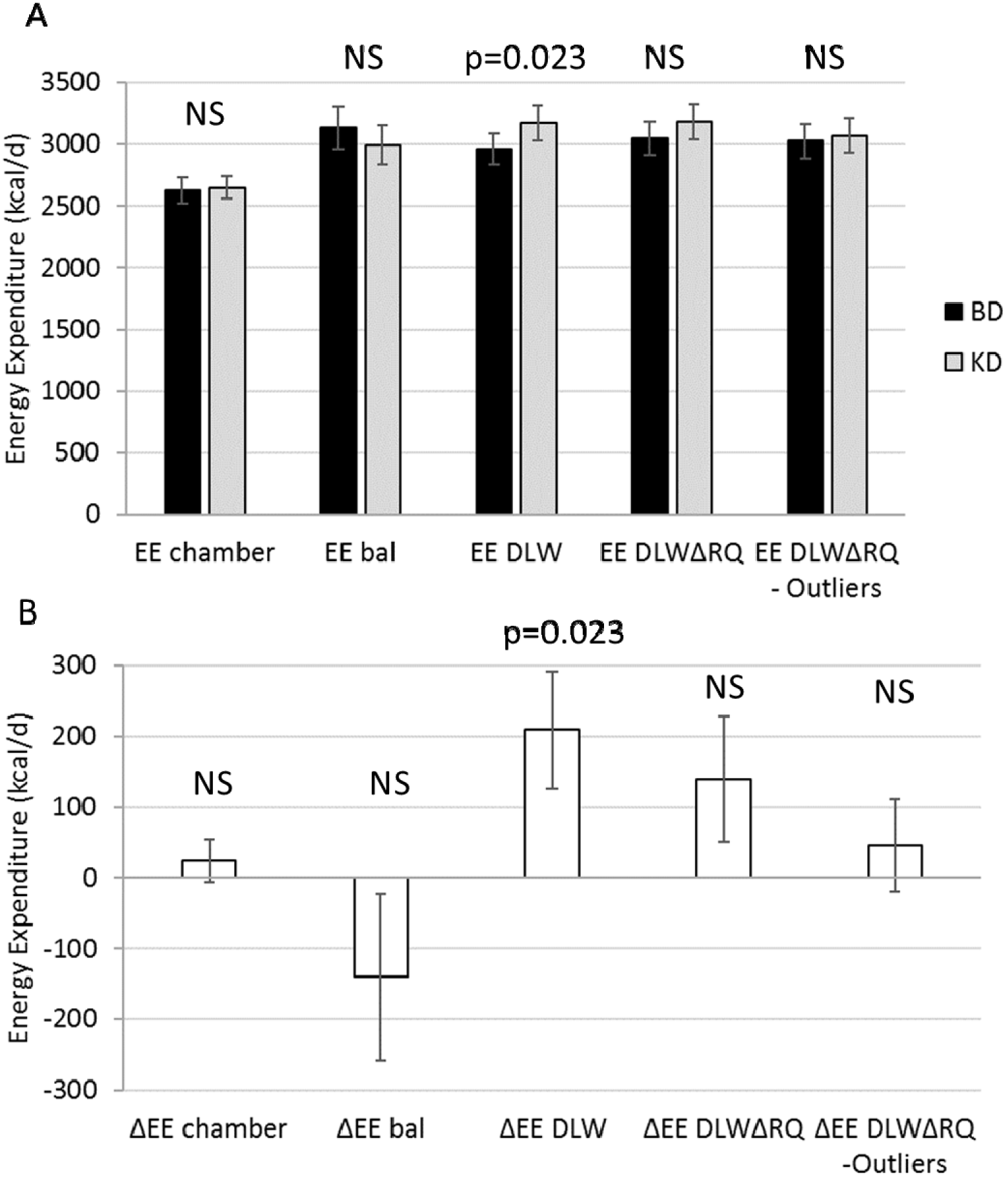
A) Neither energy expenditure by respiratory chamber (EE_chamber_), nor energy expenditure by balance (EE_bal_) were significantly different during the baseline diet (BD) versus the ketogenic diet (KD). However, energy expenditure by doubly labeled water (EE_DLW_) was significantly greater during the KD, but not after adjustment of the respiratory quotient (RQ) to account for the differential diet effect of energy imbalance (EE_DLW_ΔRQ) or after removal of two outliers whose DLW data were incomparible with other measurements (EE_DLW_ΔRQ - Outliers). B) Differences in EE_chamber_, EE_bal_, EE_DLW_, EE_DLW_ΔRQ, and EE_DLW_ΔRQ - Outliers between KD and BD phases. NS = not significant, p > 0.05.

Despite no significant differences in EI, EE_chamber_, EE_bal_, physical activity, or exercise efficiency between the KD and BD phases (Figure 1B), EE_DLW_ was 209±83 kcal/d higher during the KD phase (p=0.023). The transition from the BD to KD coincided with increases in EE_DLW_ that were in the opposite direction to EE_bal_, indicating that the DLW calculations during the KD were incommensurate with the changes in body weight and fat mass. After adjusting for the degree of energy imbalance, EE_DLW∆RQ_ was 139±89 kcal/d higher during the KD, but this difference was no longer significant (p=0.14) and was still in a direction opposite to the changes in body energy stores.

There were two clear DLW outliers. The first outlier, “subject A”, had an EE_DLW_ that was 1220 kcal/d greater than EI during the BD, and was 1751 kcal/d greater than EI during the KD despite slight body weight and fat mass gains during these periods. In contrast, the EE_chamber_ measurements for this subject were 173 kcal/d less than EI during the BD, and 65 kcal/d less than EI during the KD. The second outlier, “subject B”, had an EE_DLW_ during the BD that was only 123 kcal/d higher than EE_chamber_, but during the KD his EE_DLW_ increased by 1136 kcal/d which was ~3 standard deviations greater than the mean increase in EE_DLW,_ suggesting severe negative energy balance despite the subject gaining weight during this period and an EE_chamber_ increasing by only 72 kcal/d. Tables S1-S4 in the Online Supplemental Materials provide summary data on the energy expenditure comparisons between BD and KD phases with and without the exclusion of these subjects. After excluding these subjects, the increase in EE_DLW_ during the KD was 126±62 kcal/d (p=0.063) and EE_DLW∆RQ_ increased by only 46±65 kcal/d (p=0.49), neither of which were significant (Table S4)

## Discussion

As previously reported (9), our inpatient isocaloric feeding study showed that four days of respiratory chamber measurements that were coincident with each two-week DLW period did not detect significant changes in energy expenditure that were attributable to diet composition. However, the increase in EE_DLW_ after transitioning to the KD was substantially greater than could be accounted for by changes in body energy stores. In the present analysis, after employing diet-specific RQ adjustments in the DLW calculations to account for energy imbalance (12), we found that the calculated increase in DLW energy expenditure during the KD was significantly attenuated and was no longer statistically significant. After removal of two clear DLW outliers, the RQ-adjusted increase in DLW energy expenditure was further reduced to a marginal ~50 kcal/d higher during the KD.

We previously hypothesized that the discrepancy between the respiratory chamber and DLW measurements was due to increased physical activity outside the respiratory chamber during the KD (9). However, this potential explanation does not agree with the lack of increase in objectively measured physical activity. Also, the energy expended to perform the same low-intensity exercise in the respiratory chamber was not significantly changed by the KD, making it unlikely that the energy efficiency of skeletal muscle contraction had been altered after transitioning to the KD. Thus, the EE_DLW_ measurements were discordant with several independent measures indicating that the KD did not result in significant energy expenditure changes. The appropriately adjusted RQ values resulted in DLW data more commensurate with the other data, especially after removing the pair of outliers.

Another methodological concern with the DLW method is the theoretical possibility that CO_2_ production rates can be influenced by the fluxes through biosynthetic pathways that likely vary substantially depending on the carbohydrate content of the diet, especially the de novo lipogenesis pathway (16-18). However, the magnitude of this potential bias in humans is thought to be relatively small, amounting to an energy expenditure difference of only about 30-60 kcal/d (see Online Supplemental Materials). Interestingly, even this small systematic bias would have been sufficient to nullify the statistical significance of the observed increased EE_DLW_ in our study and the effect size was similar to the nonsignificant increase in the EE_DLW∆RQ_ difference after removal of the outliers. Individual variability of de novo lipogenesis changes after transitioning to the KD might explain the diminished correlations during the KD between the DLW measurements with EE_chamber_ and EE_bal_. To directly address this question, we need a validation study in humans with several days of simultaneous respiratory chamber and DLW measurements high-and low-carbohydrate diets, ideally with simultaneous measurements of de novo lipogenesis.

Ebbeling et al. (5) reported significant ~200-300 kcal/d differences in EE_DLW_ during the last two weeks in each of three consecutive month-long periods during which outpatient subjects consumed a very low-carbohydrate diet (10% of energy) compared to isocaloric diets with moderate (40% of energy) or high (60% of energy) carbohydrate in random order. However, resting energy expenditure differences were much smaller (~29-67 kcal/d) and objectively measured physical activity was not significantly different among the diets. Despite claims of weight stability, reported energy intake was several hundred calories below the measured EE_DLW_ but RQ was estimated without adjustment for the subjects’ state of energy balance. Therefore, similar to the current study, the diet specific effects of energy imbalance on RQ likely contributed to the observed differences in EE_DLW_. Thus, the data of Ebbeling et al. could be interpreted as supporting the possibility that the EE_DLW_ calculations using RQ values unadjusted for energy imbalance in a diet-specific way were systematically biased such that the isocaloric very low-carbohydrate diet resulted in systematically increased EE_DLW_.

In contrast, Bandini et al. (19) found in an outpatient study that EE_DLW_ was lower during the very low-carbohydrate diet (~7% of energy) as compared to a high-carbohydrate diet (~83% of energy), but this reduction was attributed to decreased physical activity because the subjects reported nausea and lethargy on the low-carbohydrate diet. No significant differences in REE were found. Stubbs et al. (20) found no significant difference between EE_DLW_ using a narrower range of diets with 29-67% of energy as carbohydrate. But the diets were fed *ad libitum* and energy intake on the low-carbohydrate diet was greater than the moderate-carbohydrate diet which was greater than the high-carbohydrate diet. Thus, the variation in total carbohydrate content of the test diets was attenuated such that daily carbohydrate intake varied by only ~22% of the mean energy intake between diets which may not have been a sufficient range to observe a systematic bias of the DLW method. Furthermore, the positive energy balance with the low-carbohydrate diet increased RQ whereas the negative energy balance with the high-carbohydrate diet decreased RQ. Thus, the RQ differences during consumption of these diets were attenuated by the differences in energy balance.

It is important to emphasize that our study was not intended to be a DLW validation study and there were several limitations. The DLW measurements were not pre-specified as either primary or secondary endpoints of the study. Whereas respiratory chamber measurements have high-precision, with an intrasubject coefficient of variation of EE_chamber_ ~2-3% (21), the DLW method is less precise, with an intrasubject coefficient of variation of energy expenditure of ~ 8-15% (22). Therefore, the relatively large inherent variability of the DLW method may have led to an apparent increase in EE_DLW_ during the KD simply by chance (type-1 error). However, we cannot definitively exclude the possibility of a real increase in energy expenditure, especially at an effect size of ~50-140 kcal/d after excluding two likely DLW outliers or using diet-specific adjustments of the DLW calculations to account for the energy imbalance.

The DLW method has been validated during 30% caloric restriction with a 55% carbohydrate diet (23) and agrees with our result that EE_DLW_ and EE_DLW∆RQ_ were not significantly different from EE_bal_ during the BD diet phase. Nevertheless, the calculated EE_bal_ values are somewhat uncertain because DXA has a limited ability to precisely and accurately detect small changes in body energy stores (24). We cannot rule out the possibility that the KD resulted in undetected increases in activity-related energy expenditure that were undetected by accelerometers. Finally, the order of the diets was not randomized, and it is possible that the elevated EE_DLW_ occurred simply because the KD followed the BD. Indeed, others have reported a greater metabolic rate during a low-carbohydrate diet when it followed a high-carbohydrate diet as compared with the reverse order (25).

In summary, our data illustrate the challenges of using the DLW method to estimate energy expenditure differences between diets varying widely in the proportion of carbohydrate. We urge caution when interpreting such data, especially if the DLW calculations do not appropriately adjust RQ for energy imbalance or if the DLW results are not corroborated by quantitatively commensurate observations of energy intake and body composition change.

## Acknowledgements

Nutrition Sciences Initiative (NuSI), convened the research team, helped to formulate the hypothesis and provided partial funding. NuSI and its scientific advisors were given the opportunity to comment on the study design and the manuscript, but the investigators retained full editorial control. The authors’ contributions were as follows: KDH, KC, RLL, MLR, MR, SRS, and ER designed the study and conducted the research; KDH and JG analyzed the data; KDH, KC, RLL, MLR, MR, SRS, and ER wrote the manuscript; KDH had primary responsibility for the final content. The authors have no conflicts of interest. We thank Jim DeLany, Herman Pontzer, and John Speakman for helpful discussions regarding the DLW method and its interpretation. The clinical study protocol and deidentified individual data are currently available for download on the Open Science Framework website at https://osf.io/h4xju/ for any purpose by anyone.

